# Information-making processes in the speaker’s brain drive human conversations

**DOI:** 10.1101/2024.08.27.609946

**Authors:** Ariel Goldstein, Haocheng Wang, Tom Sheffer, Daria Lioubashevski, Mariano Schain, Zaid Zada, Leonard Niekerken, Bobbi Aubrey, Samuel A. Nastase, Amir Taubenfeld, Harshvardhan Gazula, Colton Casto, Werner Doyle, Daniel Friedman, Sasha Devore, Patricia Dugan, Avinatan Hassidim, Yossi Matias, Orrin Devinsky, Adeen Flinker, Uri Hasson

## Abstract

Natural human conversation is driven by the exchange of information-rich messages that surprise the listener and deviate from predictable context. While extensive research has characterized how the brain processes unexpected linguistic input during comprehension, the neural mechanisms underlying the generation of such information by the speaker remain poorly understood. In this study, we hypothesize that the speaker’s brain actively creates information through extra neural computation, transforming internal thoughts into novel, meaningful linguistic outputs. Utilizing a unique 24/7 dataset of continuous electrocorticography (ECoG) comprising approximately 100 hours of spontaneous natural conversations, we contrast the neural basis of speech production and comprehension within the same participants. Behaviorally, we find that speakers take longer pausing for an additional 100–150 milliseconds before producing information-rich, improbable words, even when controlling for word identity and frequency. Neurally, we provide converging evidence for a previously unreported process: generating information-rich words elicits significantly stronger neural activity (ERP) and enhanced neural encoding and decoding in language-related areas starting 100–500 ms before articulation. This pattern contrasts sharply with speech comprehension, where enhanced neural activity for predictable words occurs before onset, while responses to improbable words emerge only after onset as prediction errors. Furthermore, we demonstrate that large language models (LLMs) mirror this biological process, requiring deeper internal computation across layers to generate improbable versus probable words. Together, these results reveal that the speaker’s brain is not merely a transmission channel but an active generator of information, recruiting extra neural resources to create novel content that diverges from listener expectations.

## Introduction

Listeners naturally prefer information-rich conversations as predictable utterances add little to what they already know. Shannon’s information theory and predictive coding frameworks help quantify the amount of information a message conveys, given its context. For example, the suffix of the statement “Look, the sky is blue” is highly predictable and therefore low in informational value, whereas the suffix of “Look, the sky is white” surprises the listener and provides new information. Classical information theory focuses on how a sender can efficiently compress a message and how a receiver can decode it. However, these frameworks have little to do with the selection of words (the message), nor do they explain the neural computations that distinguish information-rich from information-thin utterances. Psychologically, speakers aim to produce information-rich utterances to maintain their listeners’ engagement. Spontaneous speech illustrates this vividly: speakers continuously produce new, context-sensitive utterances that can surprise listeners and convey additional information. Yet, perhaps surprisingly, the neural basis of “information making” in the speaker’s brain is still one of the least studied and least understood aspects of human cognition.

Recent theoretical developments have begun to extend classical information theory by addressing not only how messages are compressed and transmitted, but also how new messages are generated. In particular, emerging frameworks propose that information can arise as a *computational product* of neural processing ^1^. Building on this idea, we hypothesize that the speaker’s brain actively creates information through neural computation, transforming internal thoughts into novel, meaningful linguistic outputs. In this view, the brain is not merely a channel for transmitting messages but an active generator of informative language. Consequently, we predicted that during free conversation, generating information-rich—as opposed to information-thin—utterances would require extra neural computation.

To explore this hypothesis, we gathered a unique 24/7 dataset of continuous electrocorticography (ECoG) for conversational analysis ^2^. The dataset comprises of spontaneous conversations throughout the patients’ day-to-week-long stays. In our setup, patients are free to generate speech and express themselves as they wish, at any time; each conversation has its unique context and purpose. Thus, for the first time, we can study the neural basis of spontaneous information making during speech production and contrast it with the process of information processing during speech comprehension within the same set of participants. This ambitious effort resulted in a uniquely large ECoG dataset of natural conversations, with approximately 50 hours (230,238 words) of neural recordings during speech production and 50 hours (289,971 words) during speech comprehension. Moreover, the superior spatiotemporal resolution and signal-to-noise ratio (SNR) of our 24/7 ECoG recordings enable us to focus this paper on the neural processes involved in generating information-rich words in the speaker’s brain and comprehending them in the listener’s brain.

In line with our hypothesis, we provide converging evidence for an additional, previously unreported neural process in the speaker’s brain that precedes the generation of information-rich (improbable) versus information-thin (probable) words within the same message. Behaviorally, speakers take longer to produce the same word when it appears in an improbable, information-rich context, indicating increased cognitive effort. Neurally, we observe that producing the same word in an information-rich context elicits enhanced ERP responses, alongside stronger neural encoding and decoding of each informative word. These complementary analyses demonstrate that the neural signals in language-related areas contain more information about upcoming information-rich words. Finally, we show that large language models also engage in additional computational steps during the generation of information-rich compared to information-thin words. Together, these results reveal that generating information-rich linguistic content recruits extra neural computation in the speaker’s brain. Such computation supports the creation of novel, meaningful content that diverges from listeners’ expectations.

## Results

### The Data Set

Our 24/7 conversation data consists of half a million words recorded during one-hundrad hours of spontaneous conversations between four ECoG patients and their surroundings in the hospital room. The conversations cover various real-life topics, including discussions between the patients and medical staff and personal conversations about family, friendships, sports, and politics. This unique dataset provided us with a rare window into the neural processes underlying the generation of information-rich utterances in the speaker’s brain during natural conversations.

To measure the amount of information communicated through words in the conversations, we used Llama-2 LLM to assess the predictability of each word within each conversational context. We operationalized word predictability using Llama-2’s predictive accuracy, conditioning on a context window of 32 words. Words successfully ranked among its top two next-word predictions were classified as *probable*. To match group size, *improbable* words were defined as those ranked below 22 in the model’s output distribution. The improbable words ranking ranged from 23 to 19019, with a mean of 319 (see Supp. Table 1). To eliminate confounding factors related to word identity, we restricted all primary results to shared words (tokens) that were classified as both probable and improbable in different contexts. This approach ensured that the observed effects were influenced by contextual information rather than by the words themselves. Similar results were observed when we included all words, as detailed in the supplementary analyses. We also replicated the results for the probability score obtained with GPT2-XL, a widely used model in previous neuroscientific research ^3–7^.

### Behavioral analysis

We show that speakers tend to slow their speech rate and pause for an additional 100–150 milliseconds before pronouncing improbable, information-rich words. This behavior aligns with the hypothesis that generating informative words requires extra neural processing (p < .001; for complete statistics, see Supp. Table 2). This pattern was observed in all four participants with neural recordings (S1–S4) and in the larger group of conversational partners whose brain activity was not recorded. Significantly, the effect was not driven by word frequency as the analysis was restricted to the subset of words shared across the probable and improbable lists (p < .001; Fig. 1). Furthermore, the same effect was seen when using all words (i.e., not just shared; see Supp. Fig. 1A), while including word frequency as a covariate (see Supp. Fig. 1B). Finally, the results are not sensitive to the model used to assess surprise, as they were replicated with the Llama-2 LLM (aee Supp. Fig. 1C).

**Figure 1.**
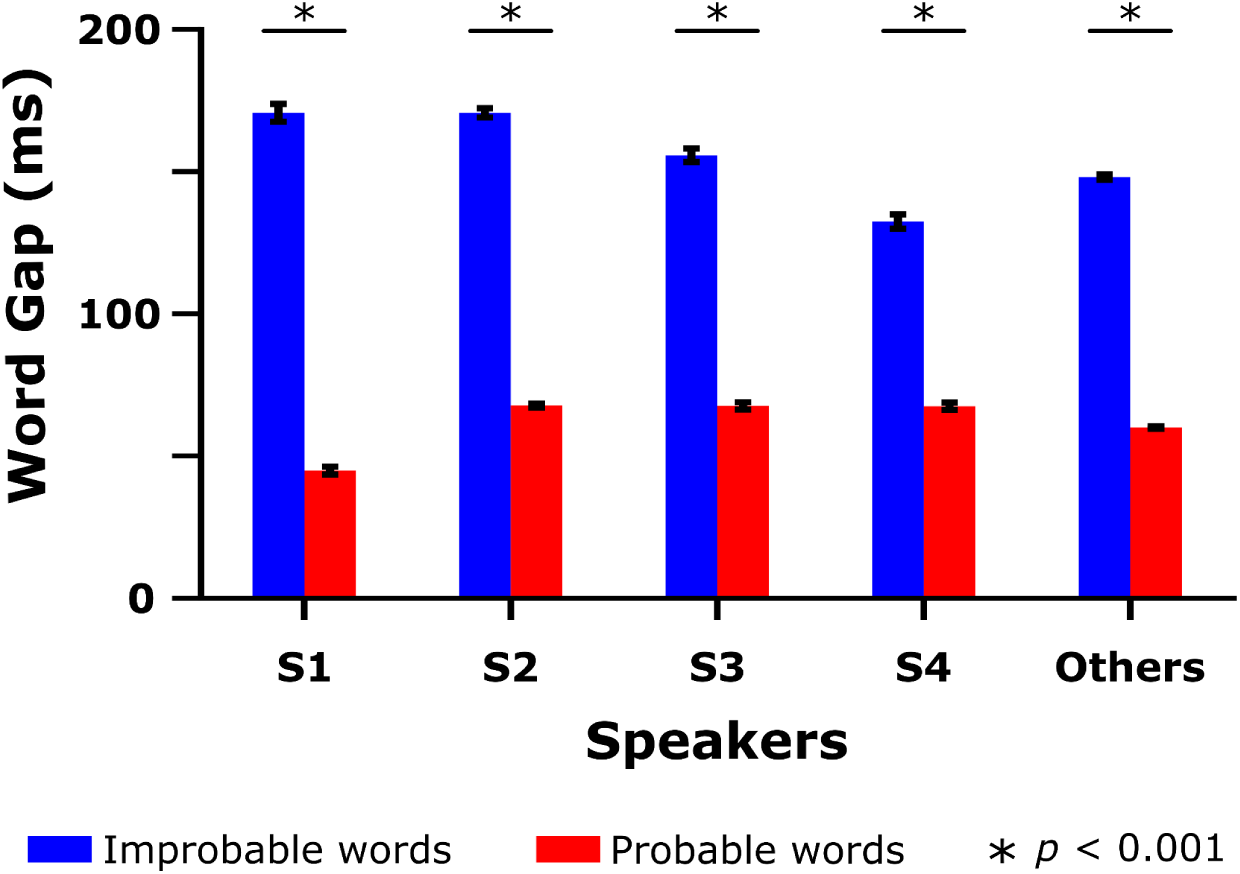
Behavior Temporal Gap between the Offset of the Previous Word and the Onset of the Current Word. It takes about 100 - 150 ms longer for each speaker (S1-S4) to start articulating improbable words. This was also evident when examining the data of other speakers in the room, for whom brain responses were not recorded. The analysis was limited to a shared set of words across both improbable and probable word lists. This suggests that the pause was independent of the word’s frequency in the natural language.

### Neural Analysis

To provide a comprehensive analysis of how the brain generates information during language production and processes information during comprehension, we examine neural responses to probable and improbable utterances using three complementary analyses. We start with a broad Event-Related Potentials (ERP) analysis of the average activity to improbable and probable spoken words, progress to word-level encoding, and finally focus on decoding both improbable and probable words from neural activity. Critically, each analysis captures different aspects of the neural responses. While ERP captures the average activity level for probable and improbable words, encoding analysis correlates specific word features with fluctuations in response on a trial-by-trial basis. This analysis is performed after removing the average neural response measured by ERPs. Consequently, encoding analysis reveals distinct, word-feature-specific aspects of the neural response that are not visible through amplitude-based analyses alone. Higher encoding signifies that a greater portion of the word representation captures the neural signal, which can be interpreted as the brain allocating more neural resources to representing or generating that specific word. Furthermore, encoding analysis employs a regression method that allows for the explicit incorporation of potential confounding variables - such as word length, inter-word gaps, and other temporal factors - as separate regressors. This approach ensures that these lower-level variables do not account for the effects attributed to semantic or lexical representations. Finally, we conduct decoding analyses, which represent the most conservative and rigorous test. In this approach, the model must differentiate between individual words based solely on their neural signatures. Successful decoding indicates that neural patterns contain enough detailed information to distinguish between improbable and probable words.

We start by analyzing signals within the inferior frontal gyrus (IFG; Fig. 2), a key area for language processing, also includes the the Broca’s area, that is involved in comprehending and producing words ^3,6,8–10^. Afterwards, we expand the analysis to other parts of the language network. In all cases, we focused on electrodes selective for language processing ^2^. This combined anatomical and functional approach guaranteed that we examined neural signals from regions both theoretically and empirically linked to language production.

**Figure 2.**
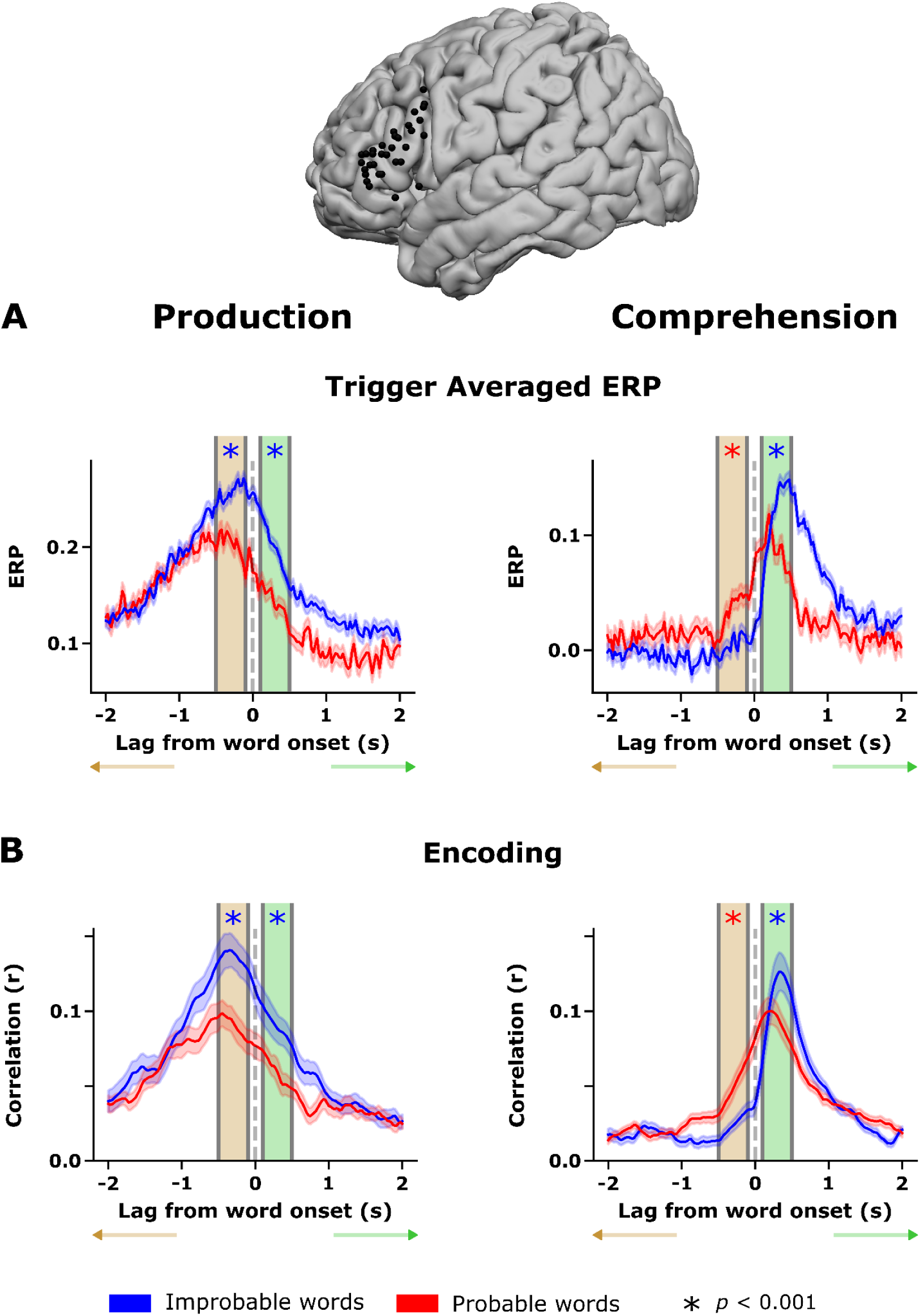
Distinct neural signatures of information generation and information processing in the inferior frontal gyrus (IFG). (A) Trigger-averaged neural responses (ERPs) in the IFG during speech production (left) and speech comprehension (right), time-locked to word onset (0 ms). During production, improbable (information-rich) words elicited significantly stronger neural activity than probable (information-thin) words beginning ∼100–500 ms before word onset, indicating enhanced preparatory neural processing prior to articulation. In contrast, during comprehension, probable words showed elevated pre-onset activity consistent with anticipatory prediction, whereas improbable words evoked stronger post-onset responses reflecting prediction-error processing. Shaded regions denote ± s.e.. (B) Word-level encoding results in the IFG using non-contextual Llama-2 embeddings. Encoding performance (correlation between predicted and observed neural responses) is shown separately for probable and improbable words across time lags relative to word onset. Consistent with ERP findings, encoding performance was higher for improbable words during pre-onset phases in production, whereas in comprehension, enhanced encoding for improbable words emerged primarily after word onset. All analyses were restricted to a shared set of words that were predictable in one context and unpredictable in another, controlling for word identity, frequency, and articulatory factors. Asterisks indicate time points with significant differences between conditions (p < .005). (B) Pre-Onset Pattern in Comprehension: In contrast to production, the listener’s brain shows a reversed pattern before word onset, with stronger encoding for probable (predictable) words. This effect is most prominent in the IFG and STG, reflecting the brain’s active engagement in predicting upcoming speech. (C)Temporal Dynamics Across Production ROIs: Analysis of separate regions of interest (ROIs) during production reveals that the enhanced encoding for improbable words is not transient; it emerges 100–500 ms before articulation and persists into the post-onset period. These sustained effects are particularly robust in the Superior Temporal Gyrus (STG) and Supramarginal Gyrus (SM). (D) Post-Onset Shift in Comprehension: Following word onset in listeners, the neural preference shifts from predictable to improbable words across the network. Of 119 speech-responsive electrodes, a significant majority (52 vs. 13) showed a preference for improbable words after onset, marking the processing of "surprisal" or prediction errors.

### Trigger-averaged neural activity (ERP)

In support of our hypothesis that generating information requires additional neural processing, we observed that during speech production, there is a stronger neural response to improbable (i.e., information-rich) words than to probable (information-thin) words, occurring 100–500 ms before the word is spoken (Fig. 2A, left). This rise in neural activity suggests that the IFG during speech production engages additional computational resources before generating information-rich utterances. To our knowledge, this is the first evidence showing an increase in neural activity around 500 ms before speech begins for information-rich utterances. The pre-onset response excludes influences from articulation or acoustic features. The rise in activity before improbable words lasted approximately 100 to 500 ms after the word started, indicating that the effect continues into early stages of articulation or immediately afterward. We remind that to dissociate information content from word duration during articulation, we restricted the analysis to a subset of shared words that were improbable in one context but predictable in another. Because the exact words appeared in both conditions, the observed differences cannot be attributed to word length, phonetic complexity, or lexical frequency. In a control analysis, we repeated the analysis for all the words.

During speech comprehension (Fig. 2A, right), we identified two distinct neural phases that closely align with the well-established distinction between pre-onset prediction and post-onset prediction-error processing ^6,11–15^. The first phase occurred before word onset, during which we observed an increase in neural activity in the listener’s brain for probable (predictable) utterances relative to improbable (unexpected) ones. This pre-word-onset increase in neural activity for predictable words is consistent with prior work showing predictive coding in the IFG ^11,12^. The second phase occurred after word onset, showing increased neural activity for improbable words relative to probable words. This increase in neural activity for unlikely words is associated with violations of listeners’ expectations (prediction-error responses) and corresponds to N400 patterns observed for surprising (i.e., less predictable) language input ^13^.

The results were replicated across multiple control analyses designed to account for potential confounding factors, including acoustic and lexical properties. First, we ran the analysis for all words (not just shared; Supp. Fig. 2A). Second, the core findings were replicated when the analysis was limited to content words only (see Supp. Fig. 2B). Third, the results were also replicated using GPT-2 predictions to classify words as probable (see Supp. Fig. 2C). Taken together, these control analyses demonstrate that the observed effects reflect information-processing driven by contextual surprisal, rather than secondary consequences of word form or articulation.

### Encoding analysis

Encoding models measure how effectively specific stimulus features (i.e., embedding) of a word can predict neural responses, helping us identify neural activity related to the properties of the words. The higher the encoding performance is, the more of the signal is explained by the word representation. Additionally, encoding models, unlike ERPs, which measure only mean amplitude differences, enable us to account for potential confounds by incorporating additional regressors. To model word-level responses, we used non-contextual word embeddings (extracted from the static embedding layer of Llama-2). We then trained encoding models (using 10-fold-cross-validation Ridge regression) for each IFG electrode for speech comprehension and speech production (separately), using time lags from –2 seconds to +2 seconds relative to word onset (lag = 0). These models were trained to predict neural responses to improbable (information-rich) and probable (information-thin) words in each conversation (see the Method section). The plot shows, at each time point, the correlation between the predicted signal and the actual neural signal.

The encoding analyses mirrored the pattern seen in the ERP results (Fig. 3). We found that encoding models predicted the neural activity for improbable (information-rich) words significantly better than for probable (information-thin) words during the pre-onset phase. This supports the claim that generating information-rich words requires additional neural processes. Specifically during the pre-onset phase, provides direct evidence that information-rich words require additional neural resources. This is because a higher encoding correlation indicates that a larger portion of the neural signal is explained by the word representation. Therefore, this increased correlation can be interpreted as the brain allocating greater neural resources toward representing or generating the word as the complexity of the information increases.

**Figure 3.**
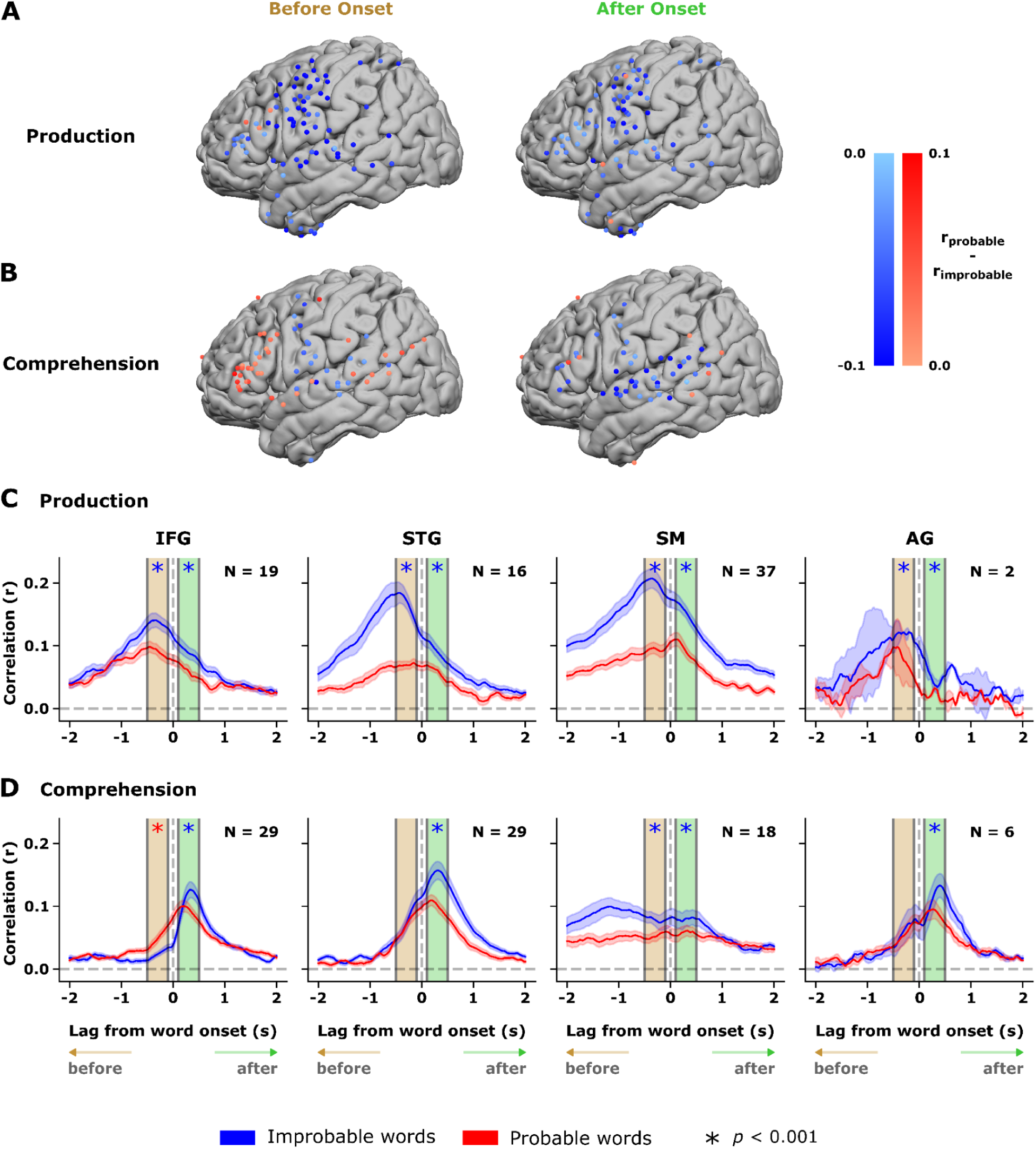
Distributed encoding of information-rich language across the cortical language network. (A) Global Enhancement in Production: During speech production, encoding is consistently stronger for improbable (information-rich) words than for probable (information-thin) words across the entire language network, including the IFG, STG, SM, and AG. This effect begins prior to word onset and remains sustained throughout articulation.

These pre-onset effects temporally precede acoustic realization and motor execution, ruling out explanations based on articulation, word duration, or auditory feedback. This timing strongly suggests that the observed neural differences reflect preparatory or planning-related processes rather than downstream motor or acoustic consequences. Notably, encoding results were obtained for shared words that were unpredictable in one context but expected in another, allowing control over word identity, length, frequency, and articulatory features. The findings remained stable when analyzing all words (see Supp. Fig. 3A), only content words (see Supp. Fig. 3B), and when defining word probability based on GPT-2 predictions (see Supp. Fig. 3C). To ensure these effects did not depend on positional information in large language model (LLM) input embeddings, we replicated all analyses using GloVe embeddings, which are pretrained without positional information (see Supp. Fig. 3D).

To rule out the possibility that pre-onset predictivity was driven by semantic similarity between consecutive words (word n and n-1), we conducted a control analysis, where we generated novel set embeddings, were the shared information between consecutive embeddings is removed. We do it by removing the projection (inner product) of each word-GloVe vector from the previous word-GloVe vector. Using these embedding, we repreduced the effect, ruling out shared information between consecutive words as confounders (see Supp. Fig. 3E). Finally, unlike the ERP analysis, the encoding framework allows multiple regressors to be included simultaneously, enabling us to explicitly model and to partial out potential confounding variables. We leveraged this flexibility to conduct an additional control analysis by regressing out word duration and inter-word pauses. This complementary encoding analysis confirmed that these temporal factors explained little, if any, variance in the neural activity (see Supp. Fig. 3F). The persistence of the pre-onset production effects under this manipulation confirms that the results are not driven by embedding correlations or lower-level acoustic and lexical properties, but rather by the neural allocation of resources toward contextual processing.

To determine whether specific participants influenced the effects, we re-ran all encoding analyses separately for each patient (see Supp. Fig. 4). In the production task, every participant showed a significant difference between improbable and probable words both before and after word onset. In comprehension, all patients exhibited the same directional pattern, higher encoding of improbable words before and after word onset. This pattern reached significance for three out of the four participants. Overall, the per-patient results closely match the group-level findings, indicating that the effects are not driven by any single participant and supporting the robustness of our main conclusions.

**Figure 4.**
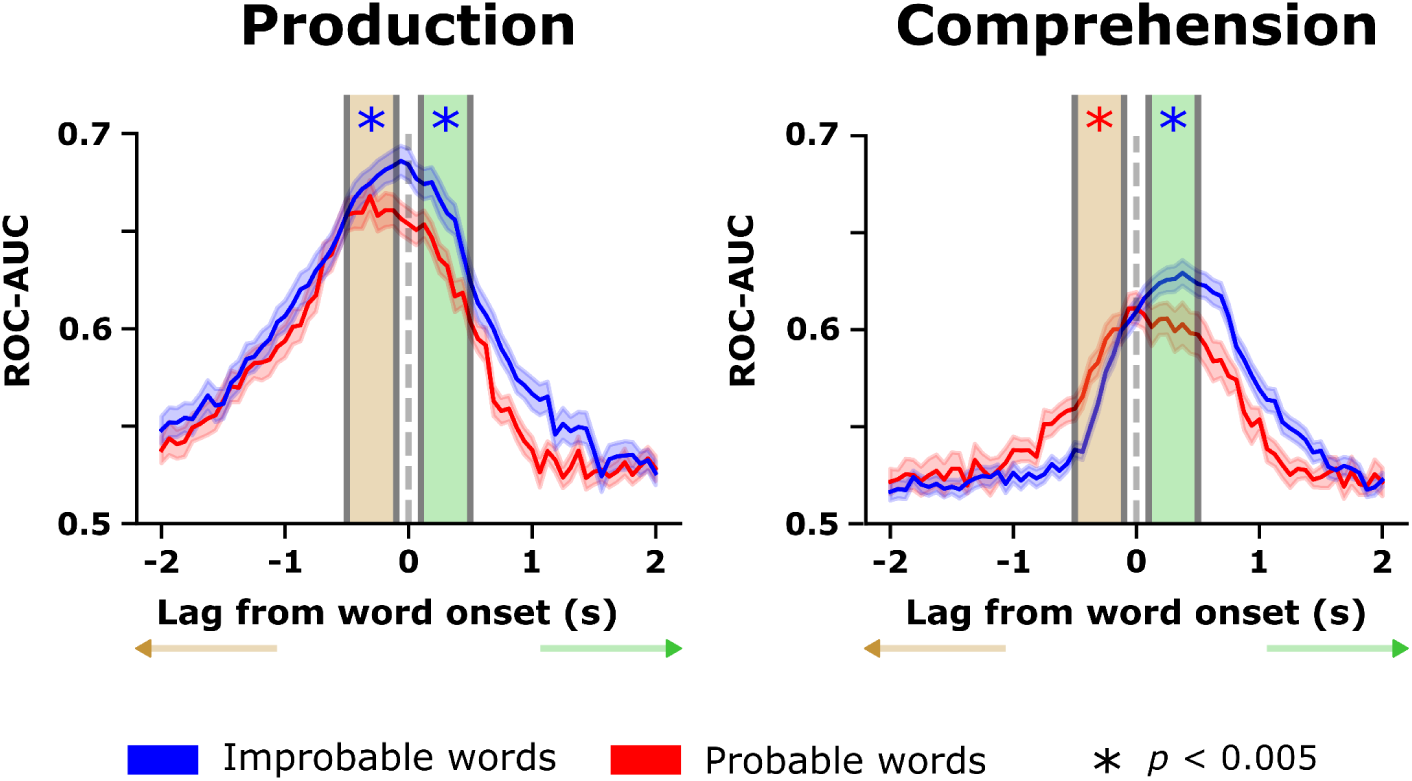
Decoding of probable and improbable words. Neural activity pooled across all language-related ROIs was used to decode individual shared words using a two-layer CNN. Decoding performance (ROC–AUC) was higher for improbable (information-rich) than for probable (information-thin) words, indicating that neural population activity carries greater discriminative information for words produced or heard in unpredictable contexts. Significance of word decoding performance was evaluated using linear mixed-effects models controlling for participant and word identity; asterisks denote significant differences (p < .005).

### Encoding analyses across the language-related ROIs

Our initial analys focuses on the inferior frontal gyrus (IFG), this choice was motivated by both theoretical and empirical considerations. The IFG is a core hub of the language network and has been repeatedly implicated in lexical selection, semantic control, and speech production planning, particularly during naturalistic language use ^8,9^. In our prior work using the same 24/7 conversational ECoG dataset ^2^, we showed that IFG plays a central role in bridging high-level linguistic representations and lower-level acoustic structure during spontaneous speech, exhibiting strong alignment with hierarchical language-model representations. For this reason, IFG provided a principled starting point for testing our hypothesis that generating information-rich language engages additional preparatory computation.

Nevertheless, language production is known to rely on a distributed cortical network, including temporal and parietal regions involved in lexical access, phonological integration, and semantic association ^2,16,17^. Our hypothesis does not predict that information making should be localized exclusively to IFG. Rather, if producing information-rich utterances requires additional neural computation, this process should manifest as coordinated changes across the broader language network. Consistent with this view, extending our encoding analyses beyond IFG revealed highly similar patterns across superior temporal, supramarginal, and angular regions. These results indicate that information generation is not confined to a single cortical locus, but instead reflects a network-level process that recruits multiple language-related areas during speech planning and articulation.

During production, encoding was consistently stronger for improbable (information-rich) words than for probable words across all examined ROIs, including IFG, superior temporal gyrus (STG), and supramarginal gyrus (SM). This enhancement emerged before word onset and persisted into the post-onset period, with particularly strong effects in STG and SM. Quantitatively, among electrodes showing significant speech-related encoding during production (123 electrodes), 110 exhibited a preference for improbable words, whereas only four preferred probable words - a highly significant asymmetry (p < .001; Fig. 3C). Enhanced encoding for improbable words also remained evident 100–500 ms after articulation, indicating sustained processing beyond word initiation. Together, these results demonstrate that generating informative language engages stronger and more readily decodable neural representations across the speaker’s language network, particularly during the preparatory phase preceding articulation.

In contrast, speech comprehension exhibited a complementary but reversed pattern (Fig. 3B). Before word onset, encoding was higher for probable words, most prominently in IFG and STG, with additional effects in SM and the angular gyrus (AG), albeit with reduced statistical power due to fewer electrodes. The pre-word-onset increase in encoding for predictable words in listeners’ IFG suggests that language areas are actively engaged in predicting upcoming words. Furthermore, post-word-onset comprehension-related encoding shifted toward improbable words in a widespread manner. Of 119 speech-responsive electrodes, 52 showed a significant preference for improbable words, compared with 13 favoring probable words (p < .001; Fig. 3D). This post-onset enhancement for improbable words shows heightened neural processing in the listener’s brain when surprising, information-rich input is received.

Taken together, these findings reveal a systematic dissociation between comprehension and production across the language network. Whereas listeners show enhanced pre-onset encoding for predictable words and post-onset sensitivity to surprising input, speakers exhibit robust, network-wide enhanced encoding for improbable words before and after articulation. This dissociation underscores the distinct computational demands of predicting versus generating language and highlights the central role of preparatory processing in the production of informative speech.

### Decoding analysis

Decoding analyses offer a more rigorous assessment of neural representations by evaluating whether neural activity provides sufficient discriminative information to distinguish individual words, rather than simply monitoring overall activation differences. Whereas encoding models evaluate how well word features predict neural responses, decoding asks whether the neural signal itself can be used to identify which word was produced or heard. Importantly, decoding leverages activity across all language-related ROIs, allowing us to aggregate unique, distributed neural information that would be lost if signals were averaged or analyzed independently. To capture these spatial patterns, we trained a two-layer convolutional neural network (CNN) with dropout, enabling the model to learn both local and distributed neural features. Decoding models were trained separately for probable and improbable words, allowing us to directly assess whether neural representations differed as a function of contextual predictability. By pooling activity across all ROIs, we aimed to decode a set of shared words that are improbable (information-rich) in one context and probable (information-thin) in another. To assess statistical significance, decoding performance (ROC–AUC) was evaluated using linear mixed-effects models with word predictability as a fixed effect and random intercepts for both participant and word identity (*auc* ∼ *prob* + (1 | *patient*) + (1 | *word*)). This approach accounts for repeated measurements across patients and lexical items while isolating the effect of word predictability on decoding accuracy. Significant effects (*p* < .005) are indicated by asterisks in the figure.

Decoding performance in the speaker’s brain was significantly higher for improbable words than for probable words before word articulation (Fig. 4A). During language production, decoding models reliably distinguished neural patterns evoked by the same words when produced in probable versus improbable contexts before word onset. In contrast, during comprehension, improved word-level decoding performance for improbable words emerged primarily after word onset, consistent with the neural processing of unexpected linguistic input (Fig. 4B). The dissociation between improved pre-onset decoding of improbable words in speech production and post-onset enhancement during speech comprehension aligns well with findings from ERP and encoding analyses. This reinforces the idea of a functional distinction between information generation in production and information processing in comprehension. We replicated the results for all words (Supp. Fig. 5).

**Figure 5.**
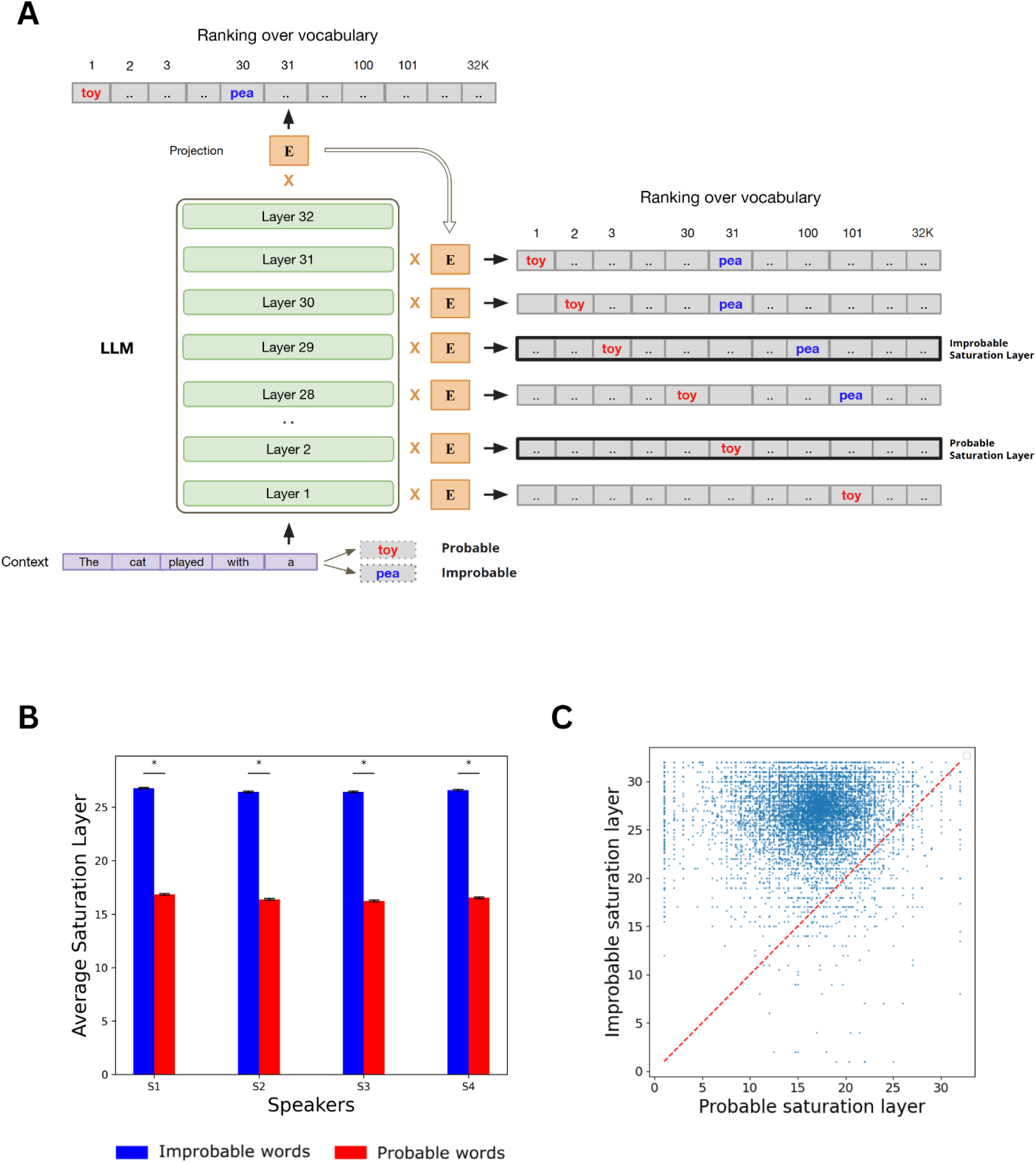
Illustration of the layer-wise saturation analysis in a language model. (A) An example illustrating how the model’s internal predictions evolve across layers. For visualization, we show two possible continuations: *toy* (probable) and *pea* (improbable), to represent a likely and an unlikely next word in this context; in the actual analyses, we track the model’s predictions for the *actual next word* and classify it as probable or improbable based on its likelihood. At each layer, the model’s hidden state is projected onto the whole vocabulary space using the unembedding matrix (E), which is typically applied only at the final layer to generate the model’s output. The resulting scores are sorted to obtain a layer-wise ranking over the vocabulary (from most to least probable). Saturation is defined as the earliest layer from which a word remains within the model’s top-100 predictions across all subsequent layers. In this example, the probable word (toy) reaches saturation early (layer 2). In contrast, the improbable word (pea) does so only much later (layer 45), demonstrating that improbable, information-rich words require deeper computation. (B) In Llama-2 (7B), improbable words reached saturation (the earliest layer after which the word consistently stays within the model’s top-100 predictions) at significantly later layers than probable words across participants (p < .001, paired t-test), even when restricted to *shared words* that appeared as both probable and improbable in different contexts. (C) A word-level analysis across subjects showed the same pattern: most words lay above the diagonal, indicating later saturation for improbable contexts. A binomial test confirmed that the proportion of words above the diagonal was significantly greater than expected by chance (p < .001).

Critically, successful decoding of shared words demonstrates that contextual predictability shapes neural representations at the population level, rather than merely modulating response magnitude. The ability to discriminate identical lexical items across contexts indicates that information-rich and information-thin words are encoded in different, distributed neural patterns across the language network. Together, these findings provide converging evidence that contextual surprisal is reflected not only in graded neural responses but also in the discriminability of neural population codes, highlighting the role of additional neural computation when generating or processing informative language.

### Computational Modeling

We next analyzed how large language models (LLMs) generate information to gain insights into the additional computation needed for generating improbable, information-rich utterances in natural language. Specifically, we input our conversations into the LLMs and asked whether information-rich words require additional internal computation to emerge as stable candidates for speech generation. Our analysis of the LLMs builds on prior work demonstrating a correspondence between the layer-by-layer accumulation of contextual information in LLMs and the temporal dynamics of neural activity in higher-order language regions ^2^.

To test our hypothesis that generating improbable, information-rich words requires additional computational resources, we examined how word predictions evolve across the layers of Llama-2. Using the logit lens ^18^ we projected hidden states from each intermediate layer into the model’s output vocabulary space via the unembedding matrix (Fig. 5A; Methods), yielding a ranked distribution over candidate next words at each layer. This allowed us to track how the rank of the actual next word changes as computation unfolds across layers. Building on prior work that characterizes saturation as the point at which a model’s predictions stabilize across layers ^19,20^, we operationalized saturation as the earliest layer at which the actual next word becomes a stable, high-ranking candidate within the model’s output space (i.e., reaches the top-100 ranked words; Fig. 5A). Specifically, saturation reflects the layer after which the true next word consistently remains among the model’s top-ranked predictions as computation unfolds. Using this measure, we quantified the depth of computation required for words to emerge as viable predictions and compared saturation across contexts for shared words.

Our analysis revealed that probable words emerged as top-ranked candidates at earlier layers than improbable words in the model, where improbable words reached saturation at significantly later layers than probable words (paired *t*-test, *p* < .001; Fig. 5B). Similar results were obtained, with fewer shared words, when we set the saturation point to top-50, top-25 (Supp. Fig. 6). A word-level analysis across all conversations revealed the same pattern: for the vast majority of shared words (see cloud of points), the average saturation layer was higher (above the diagonal) for improbable contexts than for probable contexts (binomial test, *p* < .001; Fig. 5C). The results indicate that improbable words require additional processing in LLMs to emerge as a top-ranked viable response in context. The additional computation in the LLMs for improbable words mirrors the increased neural processing observed in the speaker’s brain. This alignment suggests that the heightened neural processing during speech production may reflect a general computational principle shared by both biological and artificial systems: generating informative linguistic output requires more extensive preparatory computation than producing predictable language.

## Discussion

Natural conversation is fundamentally driven by the exchange of information that listeners do not already possess. If upcoming utterances were fully predictable, communication would add little value. In line with this intuition and with information-theoretic principles, our analyses of ∼100 hours of spontaneous conversation reveal that approximately 20–30% of spoken words are improbable and therefore information-rich. Whereas decades of research have characterized how listeners detect and process unexpected linguistic input, the neural mechanisms by which speakers generate information-rich utterances have remained largely unexplored. Here, by combining large-scale naturalistic ECoG recordings with large language models, we provide converging behavioral, neural, and computational evidence for a previously uncharacterized information-making process in the speaker’s brain that precedes the production of information-rich words.

Behaviorally, speakers consistently slowed their speech before producing improbable words, exhibiting pauses of ∼100–150 ms relative to probable words (Fig. 1). Crucially, this slowdown was not explained by word frequency, length, or articulatory complexity, as it persisted when the same lexical items appeared in predictable versus unpredictable contexts. This context-dependent delay suggests that speakers engage in additional preparatory computation when generating informative content, even though the upcoming word is already known to them. Rather than reflecting retrieval difficulty alone, this effect likely reflects additional planning or selection processes required to construct utterances that deviate from the listener’s expectations.

Neurally, this behavioral signature was accompanied by robust and systematic differences in brain activity during speech production. ERP analyses revealed enhanced pre-word-onset responses for improbable words in the speaker’s brain, providing the first evidence under natural conversational conditions that speakers recruit additional neural resources before articulating information-rich words (Fig. 2A). This pre-onset effect cannot be attributed to articulation or acoustic feedback and instead points to preparatory processes involved in linguistic generation. Encoding and decoding analyses further strengthened this conclusion. Encoding models showed stronger pre-onset predictive power for improbable words (Fig. 2B), and decoding analyses demonstrated that neural activity carried more discriminative information for improbable than for probable words before articulation (Fig. 3). Together, these findings indicate that information-rich words are more strongly represented and more readily decodable in the speaker’s brain before they are spoken.

Importantly, these production-related effects contrasted sharply with those observed during comprehension. In listeners, encoding favored probable words before word onset, consistent with predictive, information-seeking processes, and shifted toward improbable words only after word onset, reflecting prediction-error processing. This reversal between production and comprehension reveals a fundamental asymmetry: listeners optimize for prediction, whereas speakers invest additional computation in generating unpredictability. Moreover, whereas predictive effects during comprehension were relatively localized (notably in IFG), production-related effects were widespread across the language network, including IFG, STG, supramarginal gyrus, and motor-related regions (Fig. 4). This distribution suggests that information making is not confined to a single cortical locus but emerges from coordinated activity across multiple language systems.

Speech comprehension and speech production offer complementary perspectives on the relationship between information theory and LLMs in natural language processing. In speech comprehension, the primary focus is on how listeners process linguistic information from real-world linguistic input. Recent evidence suggests that, like LLMs, humans rely on a next-word predictive coding framework to compress linguistic information into an embedding space ^4,11,21,22^. Thus, in alignment with information theory, listeners constantly seek to detect information-rich words to adjust their linguistic model based on their prediction errors.

To further probe the computational nature of information making, we turned to large language models as mechanistic analogues of word generation (Fig. 5). We showed that improbable words require deeper internal computation across layers before stabilizing as viable predictions. This mirrors the increased neural processing observed in the speaker’s brain before producing information-rich words. The convergence between human neural dynamics and LLM internal computation suggests a shared computational principle: generating informative language requires deeper processing than producing predictable continuations.

A substantial body of prior work has shown that speakers modulate speech rate, pausing, and prosody as a function of word predictability, a pattern that has been successfully framed within the Uniform Information Density (UID) theory ^23–27^. UID provides a principled account of how speakers redistribute information over time to optimize transmission through a noisy channel, assuming a fixed message content. Our findings are consistent with these behavioral observations but point to a complementary computational stage that precedes them. By focusing on neural activity before word onset and by restricting analyses to identical lexical items appearing in predictable and unpredictable contexts, we isolate a process that cannot be explained by temporal redistribution of information alone. Instead, we observe enhanced neural computation in the speaker’s brain prior to producing information-rich words, suggesting that additional processing is engaged during the *generation* of informative content itself. In this view, UID captures how information is shaped for transmission, whereas our results reveal a previously uncharacterized neural process that supports information making constructing utterances that intentionally deviate from listeners’ expectations. Together, these perspectives suggest that efficient communication relies not only on managing information density during articulation, but also on preparatory neural mechanisms that enable speakers to generate novel, informative linguistic content in the first place.

Recent work has substantially advanced our understanding of the neural dynamics supporting language production and comprehension during natural communication, including the distributed cortical organization of conversation, hierarchical neural dynamics preceding speech, and the decoding of speech and articulatory signals from neural activity ^28–30^. Collectively, these studies have characterized how language unfolds in the brain across time, space, and representational levels. However, they have largely focused on general production dynamics or speech execution, rather than on how speakers actively generate *informative* content. In particular, the preparatory neural computations that distinguish information-rich from information-thin language in naturalistic speech have remained unaddressed. In contrast, our study isolates a previously uncharacterized information-making process in the speaker’s brain by combining large-scale naturalistic ECoG recordings with large language models. We show that producing unpredictable, information-rich words engages additional neural computation before articulation, and that this effect mirrors deeper internal computation in artificial language models. Together, these findings extend prior work by revealing a neural signature of information generation itself, rather than of language production more broadly.

These findings expose a conceptual gap in using classical information theory as a framework for cognitive processing. While entropy and surprisal explain how information is encoded and transmitted, they do not address how speakers intentionally generate the informative messages they wish to share with their audience. Our results point to the fact that information theory focuses on the transfer of messages and does not adress the a computation that underlies the generation in the speaker’s brain.

Several limitations remain. Although we demonstrate enhanced neural processing of information-rich words, the precise computational policy governing their selection remains unknown. Furthermore, our estimates of word predictability rely on LLMs that lack access to speakers’ shared history and private knowledge, likely leading to conservative estimates of surprisal. Importantly, such limitations would reduce, rather than inflate, observed differences between conditions, strengthening confidence in the reported effects. Lastly, while our analyses focused on fluent segments of spontaneous speech, future research should examine whether disfluencies, points of hesitation, correction, or repair, reflect transient disruptions in the same information-making processes identified here.

In summary, by leveraging naturalistic neural recordings and computational models, this study identifies a previously unrecognized information-making process in the speaker’s brain. These results complement established theories of information seeking in comprehension and suggest that generating informative language engages distinct and computationally demanding neural mechanisms. Understanding these mechanisms may be essential for explaining how humans use language to think, innovate, and communicate new ideas, and for guiding the development of artificial systems capable of doing the same.

## Methods

### Preprocessing the speech recordings

We developed a semi-automated pipeline for preprocessing datasets consisting of four main steps. First, we de-identified speech recordings by manually censoring sensitive information to comply with HIPAA regulations. Second, we used a human-in-the-loop approach with Mechanical Turk transcribers to transcribe the noisy, multi-speaker audio accurately. Third, we aligned text transcripts with audio recordings using the Penn Forced Aligner and manual adjustments for precise word-level timestamps. Finally, we synchronized speech with neural activity by recording audio from ECoG channels, achieving 20-millisecond alignment accuracy between neural signals and conversational transcripts. For a full description of the procedure, see ^22^.

### Preprocessing the ECoG recordings

We developed a semi-automated analysis pipeline to identify and remove corrupted data segments (e.g., due to seizures or loose wires) and mitigate other noise sources using FFT, ICA, and de-spiking methods ^31^. Neural signals were bandpassed (75–200 Hz), and power envelopes were computed as proxies for local neural firing rates ^32^. The signals were z-scored, smoothed with a 50-ms Hamming kernel, and trimmed to avoid edge effects. Custom MATLAB 2019a (MathWorks) preprocessing scripts were used for these steps. For a full description of the procedure, see ^22^.

### Prediction and embedding extraction

We extracted contextualized predictions and static word embeddings from GPT-2 (gpt2-xl, 48 layers) and Llama-2 (Llama-2-7b, 32 layers). We used the pre-trained model implemented in the Hugging Face library ^33^. We first converted the words from the raw transcript (including punctuation and capitalization) into whole- or subword tokens. We used a sliding window of 32 tokens (results were also replicated with 1024 tokens), advancing by one token to extract the embedding for the final token in the sequence. By encoding these tokens as integer labels, we fed them into the model and received the activations at each layer of the network (also known as hidden states). For the predictions, we extracted the model’s logits for the second-to-last token, which the model used to predict the last token. For the embeddings, we extracted the activations for the final token in the sequence from the 0th layer of the model before any attention modules. To divide tokenized words into multiple tokens, we use the prediction values of the first token and average the embeddings across the tokens. With embeddings for each word in the raw transcript, we aligned this list with our spoken-word transcript, which lacked punctuation, retaining only whole words.

### Electrode-wise encoding

We used linear regression to estimate encoding models for each electrode and lag relative to word onset, mapping our static embeddings onto neural activity. The neural signal was averaged across a 200-ms window at each lag (25-ms increments). Using 10-fold cross-validation, we trained models to predict neural signal magnitudes from GPT-2 or Llama-2 embeddings. Embeddings were standardized and reduced to 200 dimensions via PCA (we replicated results using PCA to 50 dimensions and ridge regression). Regression weights were estimated by ordinary least squares and applied to the test set. Pearson correlation assessed model performance across 161 lags from -2,000 to 2,000 ms in 25-ms increments. For a full description of the procedure, see ^11^.

### Electrode selection

A randomization method was employed to determine significant electrodes that were selective for semantic information. Each iteration involved randomly shifting embeddings (GloVe) assigned to predicted signals, breaking their connection with brain signals while maintaining their order without rolling over within the context window. The encoding procedure was then conducted for each electrode using the misaligned words, repeated 1,000 times. The score for each electrode was calculated by the range between the maximum and minimum values across 161 lags. From these, the highest value for each patient across all electrodes was recorded, forming a distribution of 1,000 maximum values per patient. The significance of electrodes was assessed by comparing the original encoding model’s range to this distribution, calculating a p-value for each electrode. This tested the hypothesis of no systematic relationship between brain signals and word embeddings, yielding family-wise error rate-corrected p-values. Electrodes with p-values under .01 were deemed significant. For a full description of the procedure, see ^11^.

### Significance test for encoding difference at the ROI level

To test for significant differences in encoding performance between probable and improbable word conditions in 17 given lags (-500 ms to -100 ms) for a specific ROI, we used a paired-sample permutation procedure: in each permutation, we randomly shuffled the labels (probable/improbable) of all observations (correlation encoding) for both conditions, and we computed that difference of the averages. A *p*-value was computed as the percentile of the non-permuted difference between the averaged correlation values for the probable and improbable words over the electrodes and lags relative to the null distribution. *P*-values less than .0005 (significance of .001 for the two-sided test) were considered significant. We used a similar paired-sample permutation procedure to test for significance for specific electrodes with samples from the 17 given lags. FDR correction was applied to correct for multiple electrodes.

### Saturation definition for computational modeling analysis

Saturation has previously been defined as the earliest layer from which the model’s top-1 prediction remains fixed across all subsequent layers ^19^, and later extended to top-k predictions ^20^. In the present work, we further generalize this notion to capture the depth of computation required for the true next word to emerge as a stable and viable candidate for generation. This extension is necessary to include improbable words, which by definition are low-ranked by the model and would often fail to saturate under earlier definitions. We therefore define saturation as the earliest layer at which the true next word enters the model’s top-k predictions and remains within this set across all subsequent layers. We evaluated this criterion for k = 25, 50, and 100. For each word and context, the saturation layer was computed as the minimum layer index satisfying this stability condition, enabling consistent estimation of saturation for both probable and improbable words.

## Supplementary Figures

**Supplementary Figure 1.**
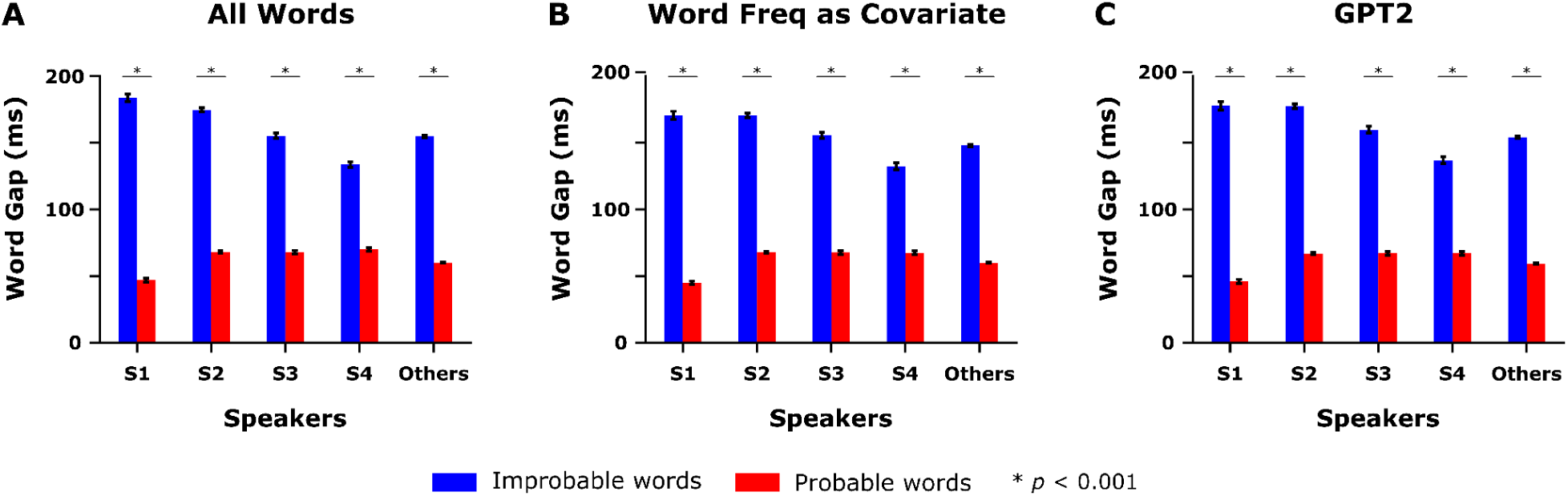
Mean word gaps across five speaker categories, comparing the temporal delay following improbable versus probable words. (A) All Words – Displays word gaps (ms) for improbable vs. probable words across different speakers (S1, S2, S3, S4, and Others). (B) Word Freq as Covariate – Shows the same effect when controlling for word frequency. (C) GPT2 – Shows the results when using the GPT2-XL LLM to assess surprise.

**Supplementary Figure 2.**
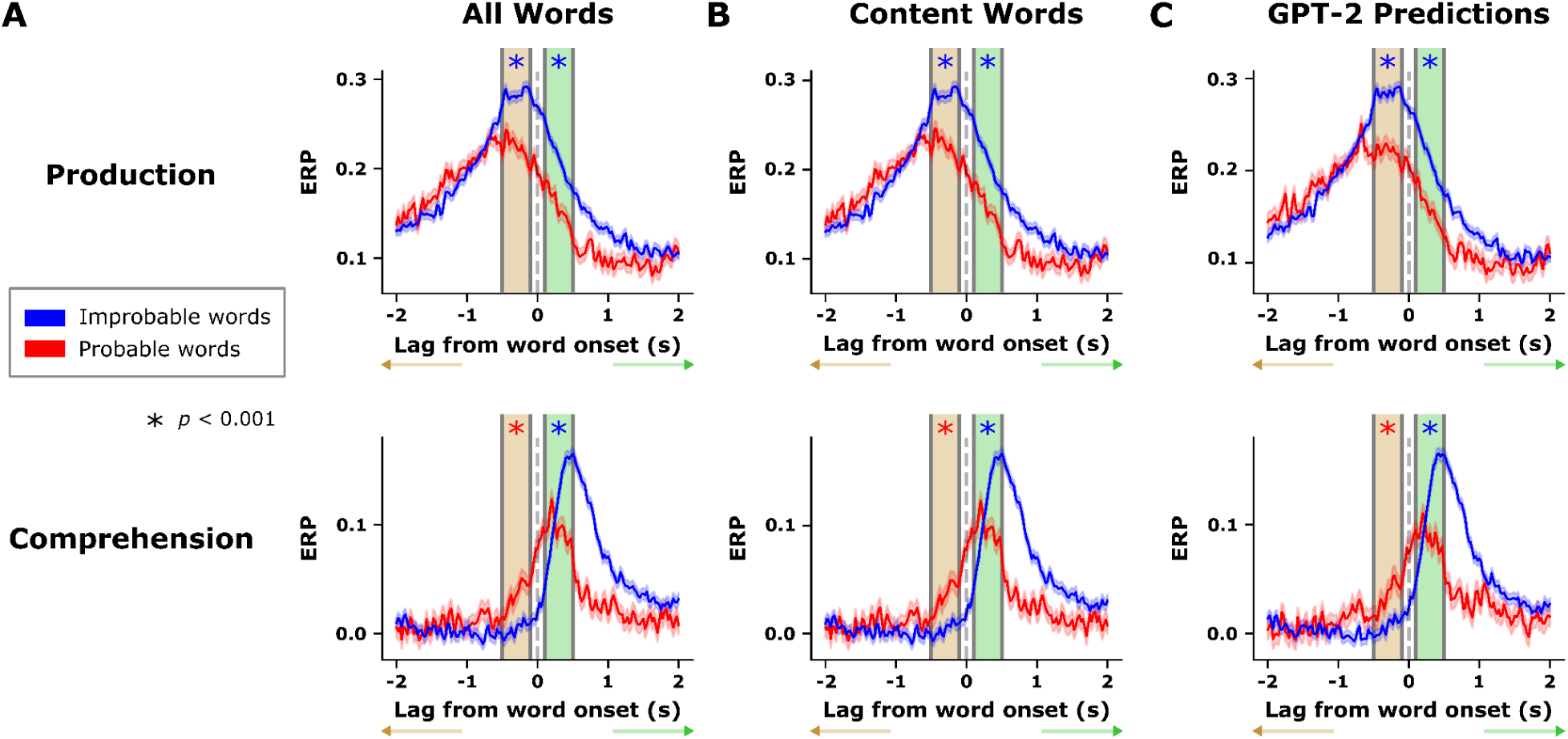
Event-related potentials (ERP) for production and comprehension tasks across various control analyses to assess the impact of contextual surprisal. (A) All Words: Analysis conducted using all words, demonstrating consistent ERP differences between improbable and probable words in both production (top) and comprehension (bottom) phases. (B) Content Words: Results replicated when the analysis was restricted specifically to content words to control for lexical properties. (C) GPT-2 Predictions: Results replicated when word probability was determined by GPT-2 model predictions to ensure findings were not sensitive to the specific model used for surprisal.

**Supplementary Figure 3.**
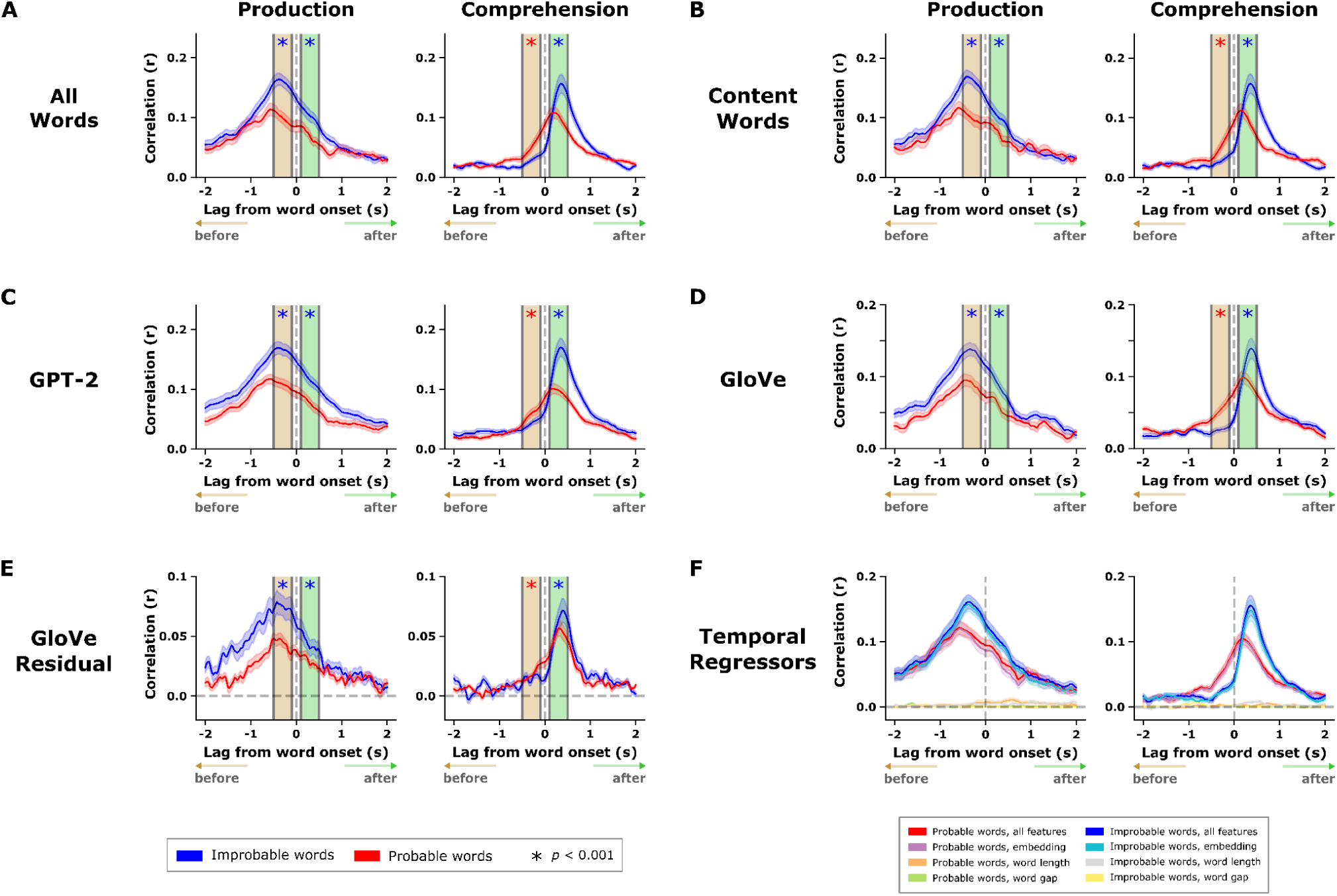
Control analyses for pre-onset encoding effects. (A) Encoding results for all words. (B) Results restricted to content words only. (C) Word probability defined using GPT-2 predictions. (D) Replication using GloVe embeddings, which are pretrained without positional information. (E) Control for shared semantic information between consecutive words. (F) Control for temporal acoustic variables.

**Supplementary Figure 4.**
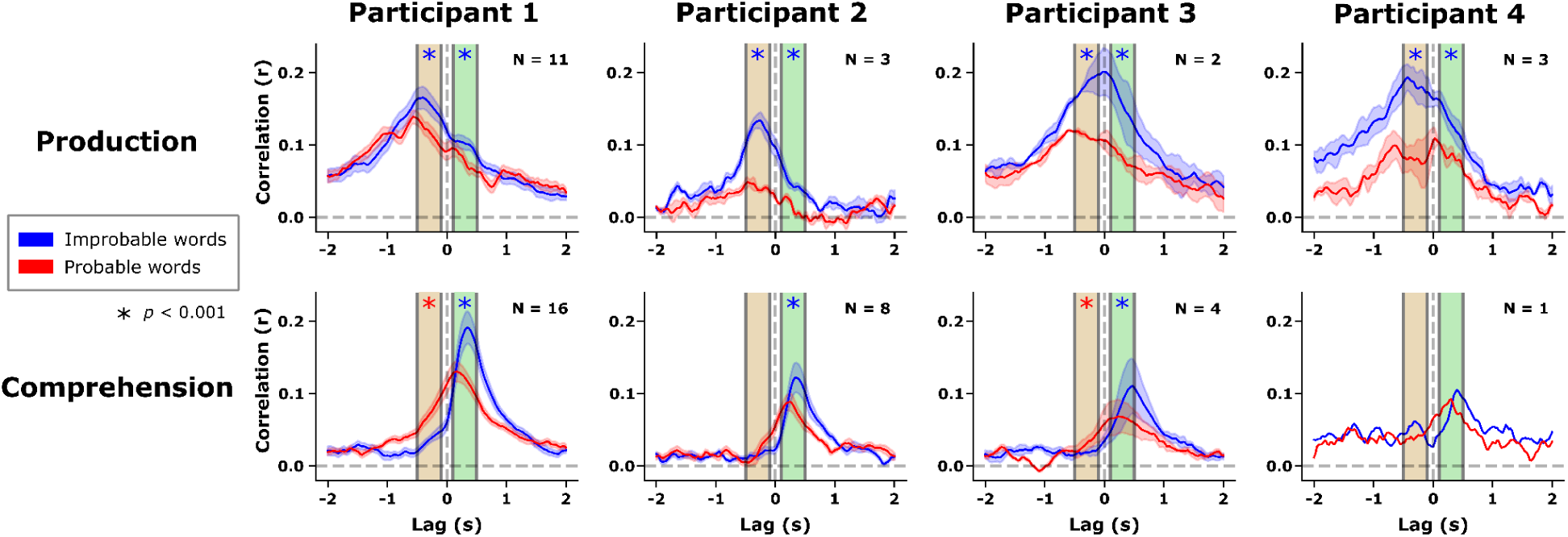
Individual participant encoding analyses demonstrate the robust nature of the observed effects across both production and comprehension tasks. To ensure that group-level findings were not disproportionately influenced by specific individuals, all encoding analyses were re-run separately for each patient. In the production task, every participant showed a significant difference in correlation between improbable and probable words both before and after word onset. During comprehension, all patients exhibited the same directional pattern—higher encoding of improbable words around the time of word onset—which reached statistical significance for three out of the four participants. Overall, these per-patient results closely match the group-level findings, confirming that the information-processing effects driven by contextual surprisal are not the result of outliers but are consistent across the sample.

**Supplementary Figure 5.**
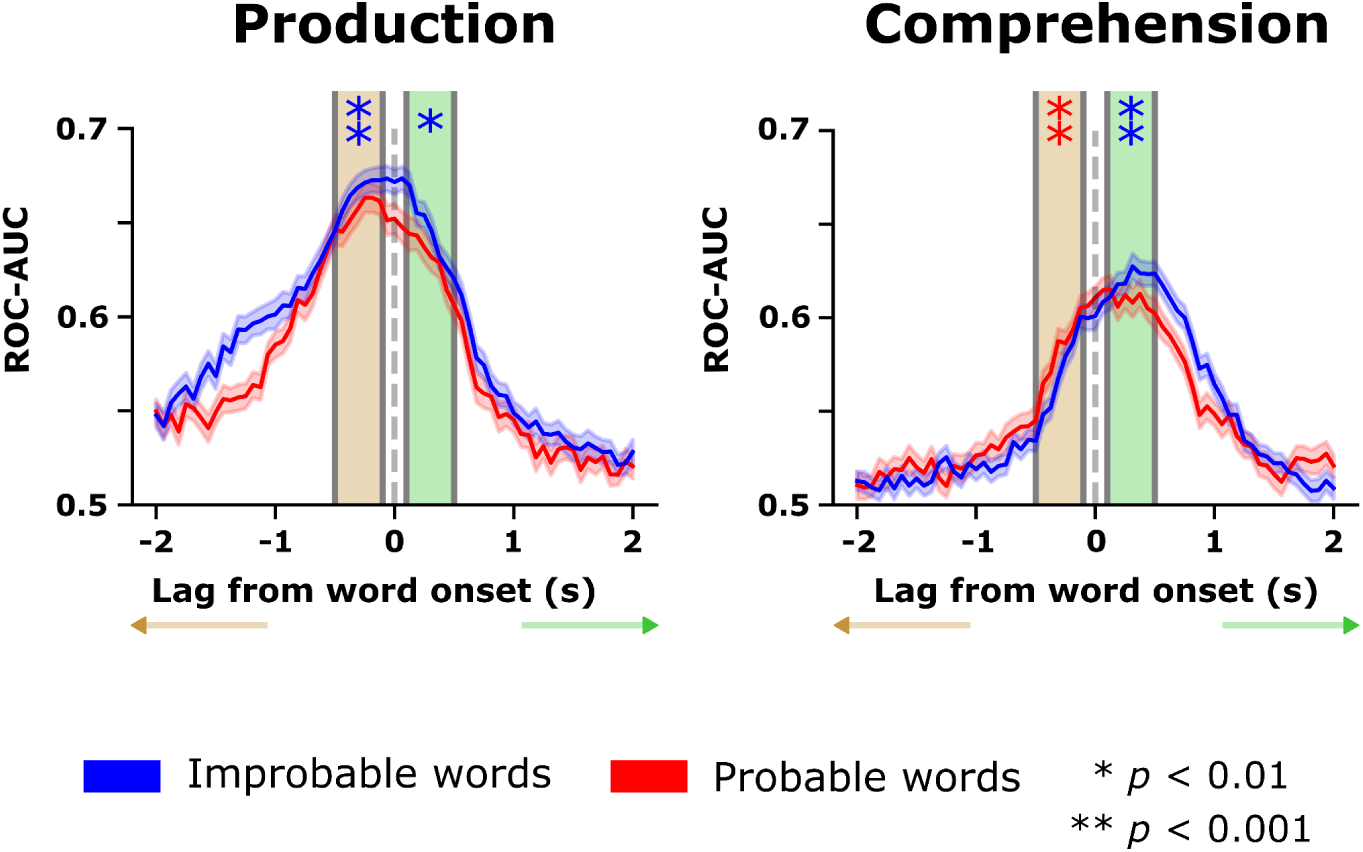
Decoding accuracy for word information using all words during speech production and comprehension. This analysis replicates the core decoding results using the full dataset of all words to demonstrate the robustness of the temporal dissociation between tasks. In speech production, decoding performance (ROC-AUC) for improbable words is significantly enhanced during the pre-onset window compared to probable words. In contrast, speech comprehension shows a post-onset enhancement, where the decoding of improbable words peaks significantly after the word has been heard.

**Supplementary Figure 6.**
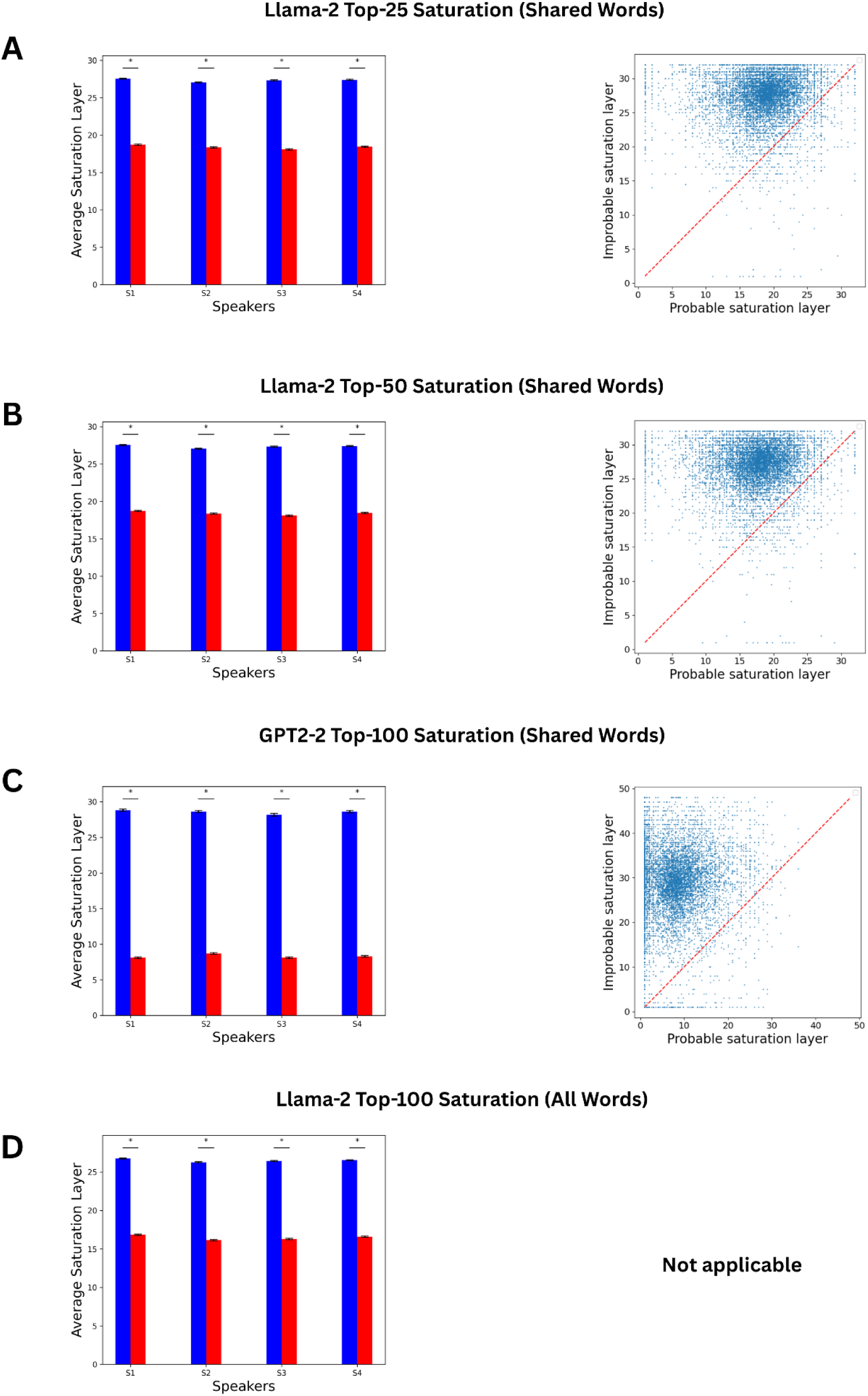
Layer-wise saturation analysis extended across different models and top-k settings. (A) Llama-2 results where saturation is defined as the earliest layer from which a word remains within the model’s *top-25* predictions across all subsequent layers. Left shows the earliest saturation layer for probable versus improbable words across participants, and right shows the word-level comparison; the pattern of later saturation for improbable words is preserved. (B) Llama-2 results where saturation is defined as the earliest layer from which a word remains within the model’s *top-50* predictions across all subsequent layers, showing the same trends as in A. (C) GPT-2 XL results (top-100 predictions) replicate the main findings: improbable words reach saturation later than probable words both across participants and at the word level. (D) Llama-2 results for all words, not restricted to shared words, using top-100 predictions; the pattern of later saturation for improbable words across participants remains consistent. In this case the word level comparison is not applicable, since words which are not shared (i.e. do not appear as both probable and improbable in different contexts) can not be placed on these axes. Across all panels, these analyses confirm that the main effect of later saturation for improbable, information-rich words is robust to model choice, top-k cutoff, and inclusion criteria.

## Supplementary Tables

**Supplementary Table 1.** Statistics of Words Divided into Probable, Improbable, and Middle (middle 40%) using Llama-2’s prediction accuracy (top table) and confidence levels (bottom table). Rank is the ranked prediction order of the next word, ranging from 1 to 32,000 (vocab size for Llama-2). Pred is the prediction probability of the next word, ranging from 0 to 1.

| Llama-2's Prediction Accuracy |  |  |  |  |  |
| --- | --- | --- | --- | --- | --- |
| Type | Word Num | Rank Mean | Rank Std | Rank Min | Rank Max |
| Probable | 173358 | 1.271 | 0.444 | 1 | 2 |
| Middle | 175626 | 8.708 | 5.393 | 3 | 22 |
| Improbable | 153661 | 318.785 | 782.165 | 23 | 19010 |

**Supplementary Table 1. Statistics of Words Divided into Probable , Improbable, and Middle (middle 40%) using Llama-2's prediction accuracy (top table) and confidence levels (bottom table).**
| Llama-2's Confidence Level |  |  |  |  |  |
| --- | --- | --- | --- | --- | --- |
| Type | Word Num | Pred Mean | Pred Std | Pred Min | Pred Max |
| Probable | 150795 | 0.420 | 0.271 | 0.084 | 0.999 |
| Middle | 201055 | 0.038 | 0.029 | 0.005 | 0.141 |
| Improbable | 150795 | $1.624e^{-3}$ | $1.691e^{-3}$ | $9.140e^{-9}$ | $7.410e^{-3}$ |

**Supplementary Table 2.**
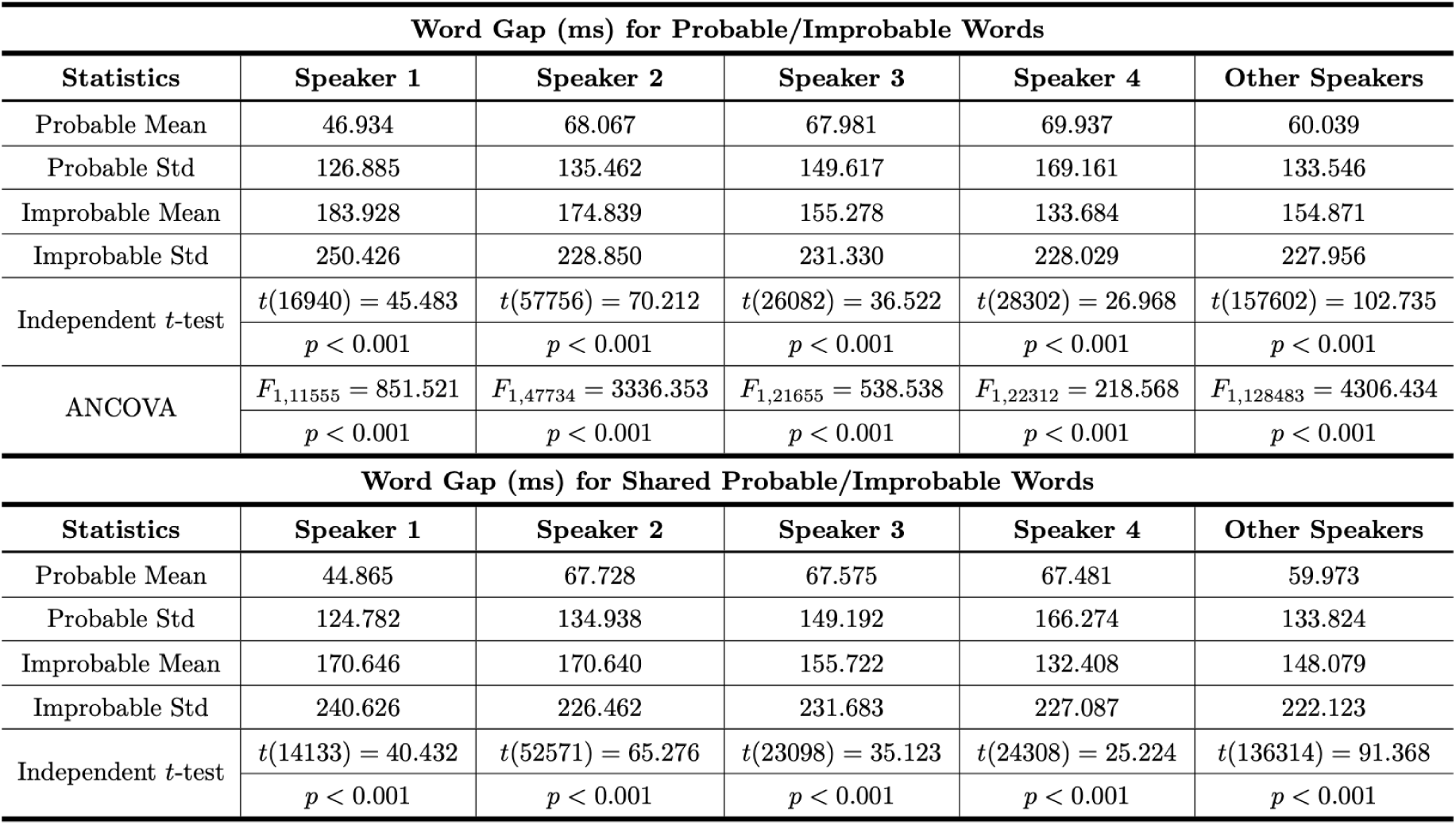
Statistics and Significance Tests for Word Gap (Duration between the offset of the previous word and onset of the current word) for probable and improbable Words.

